# Targeted Protein Degradation by KLHDC2 Ligands Identified by High Throughput Screening

**DOI:** 10.1101/2025.03.31.646306

**Authors:** Han Zhou, Tongliang Zhou, Wenli Yu, Liping Liu, Yeonjin Ko, Kristen A. Johnson, Ian A. Wilson, Peter G. Schultz, Michael J. Bollong

**Affiliations:** Department of Chemistry, Scripps Research, 10550 North Torrey Pines Road, La Jolla, California 92037, United States; Department of Integrative Structural and Computational Biology, Scripps Research, 10550 North Torrey Pines Road, La Jolla, California 92037, United States; Chemical & Biological Integrative Research Center, Korea Institute of Science and Technology, 5 Hwarang-ro 14-gil, Seongbuk-gu, Seoul 02792, Republic of Korea; Department of Biology, Calibr-Skaggs at Scripps Research, 11119 North Torrey Pines Rd, La Jolla, California 92037, United States

## Abstract

Proteolysis targeting chimeras (PROTACs) enable the selective and sub-stoichiometric elimination of pathological proteins, yet only two E3 ligases are routinely used for this purpose. Here, we expand the repertoire of PROTAC compatible E3 ligases by identifying a novel small molecule scaffold targeting the ubiquitin E3 ligase KLHDC2 using a fluorescence polarization-based high throughput screen. We highlight the utility of this ligand with the synthesis of PROTACs capable of potently degrading BRD4 in cells. This work affords additional chemical matter for targeting KLHDC2 and suggests a practical approach for identifying novel E3 binders by high throughput screening.

## Introduction

Targeted protein degradation has emerged as an alternative strategy to modulate cellular signaling. Among established pharmacological approaches, proteolysis targeting chimeras (PROTACs) have gained prominence in the field, enabling the potent degradation of several challenging cancer targets like Estrogen Receptor (ER), Androgen Receptor (AR), among others, with several progressing to ongoing clinical trials. ^1-3^ PROTACs are heterobifunctional molecules composed of a ligand targeting the protein of interest, a ligand for a ubiquitin E3 ligase, and a linker with precise geometry to bring the two proteins within sufficient proximity to induce the ubiquitination and degradation of so called ‘neosubstrate’ proteins.^4^ Despite the widespread clinical and academic use of PROTACs, most degrader molecules developed to date have exploited only two E3 ligases, VHL (von Hippel-Lindau) and CRBN (Cereblon).^5,6^ While several additional E3 ligases have been reported in the literature as being PROTAC compatible, these ligands do not have well understood mechanisms of E3 engagement and often do not fully degrade their target substrates.^7^ As such, an expanded repertoire of E3 ligands would likely enable the degradation of a broader array of protein targets to endow subcellular and/or tissue specificity of degradation or to overcome potential resistance mutations in currently targeted E3s.

Among potential E3 ligases for PROTAC development, KLHDC2 (Kelch-like homology domain-containing protein 2) has emerged as a promising candidate.^1^ A CUL2 adaptor protein, KLHDC2 serves as a key component of the C-end degron pathway, recognizing C-terminal diglycine residues of client proteins, promoting their ubiquitination and degradation.^8^ Client proteins of KLHDC2 include selenocysteine containing proteins, like SelK (Selenoprotein K), which, in the context of selenocysteine depletion, undergo premature translational termination, revealing a C-terminal diglycine degron.^9^ Previous work has shown that recognition of the diglycine degron of SelK by KLHDC2 in its Kelch beta propellor domain is characterized by low nanomolar dissociation constants (<10 nM) and with favorable binding kinetics for PROTAC development (Figure 1A).^9^ Unbiased protein proximity-based screens for generalizable E3 degraders have also indicated that KLHDC2 is among the best E3 ligases for fully degrading generic protein substrates.^10,11^ Similarly, others have shown that peptide-based PROTACs derived from a 7-mer peptide of the C-terminus of SelK can be used to degrade a variety of kinase targets in human cells.^12^

**Figure 1.**
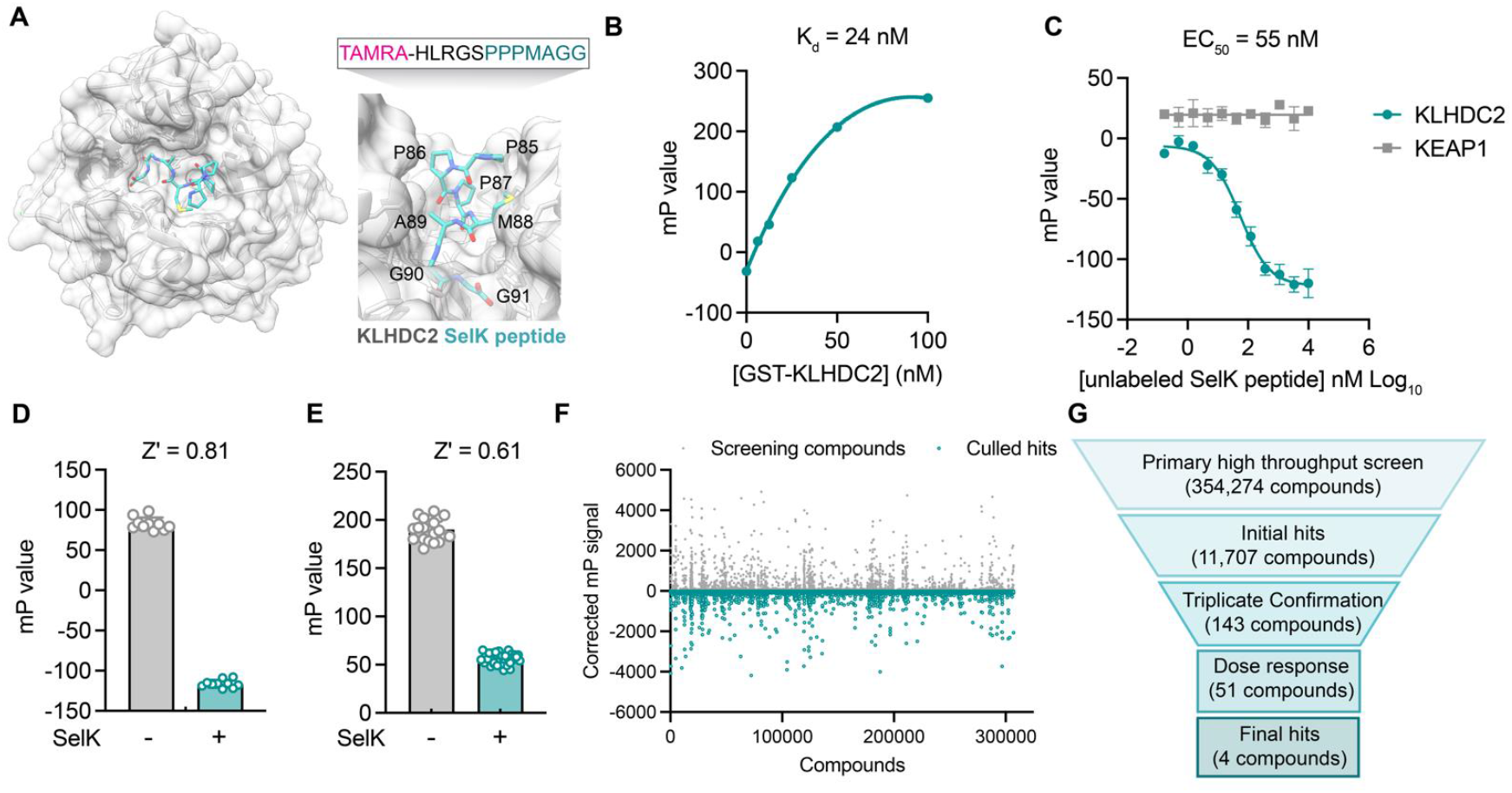
A fluorescence polarization-based screen identifies ligands of the Kelch domain of KLHCD2. A) Structure of the Kelch domain of KLHDC2 from PDB 6DO3 bound to PPPMAGG, C terminal peptide from SelK (left). Interface of SelK peptide (teal) bound to KLHDC2 (right) with TAMRA labeled peptide used in this work above. B) Fluorescence polarization signal of TAMRA-SelK peptide in response to increasing concentrations of GST-KLHDC2. C) Fluorescence polarization signal from KLHDC2 and KEAP1 assays in response to increasing concentrations of unlabeled SelK peptide. Fluorescence polarization signals in Z’ based determination assays with and without unlabeled SelK peptide (1 µM) in 384-well (D) and 1536-well (E) assays. F) Corrected fluorescence polarization signal from the primary screening campaign with hits noted in teal. G) Screening funnel depicting the high throughput screening campaign.

We reasoned that the well-characterized interaction between KLHDC2 and SelK could be exploited to identify small molecule binders of KLHDC2 from a competitive in vitro screen using recombinant protein. Accordingly, we report here the development and execution of a fluorescence polarization-based high throughput screen that identified a tetrahydroquinoline-based scaffold that competes for SelK peptide binding. We demonstrate this scaffold can be optimized for potency, achieving submicromolar affinity for KLHDC2 binding and can be further derivatized with JQ1 to degrade BRD4 in cell-based assays.

## Results and Discussion

To identify small molecules capable of binding the Kelch domain of KLHDC2, we first established a miniaturized fluorescence polarization (FP)-based assay reporting on the KLHDC2-SelK interaction. This assay derives from literature precedent in which a C-terminal peptidic fragment of SelK (HLRGSPPPMAGG) has been shown to potently associate with purified GST tagged KLHDC2 in a recombinant AlphaScreen assay.^9^ Accordingly, we evaluated fluorescence polarization signal in response to increasing concentrations of recombinantly produced GST tagged KLHDC2 from Sf9 insect cells in the presence of a 12-mer SelK peptide N-terminally conjugated with a TAMRA fluorophore (3.1 nM, TAMRA-HLRGSPPPMAGG). Under these conditions, a slightly shifted dissociation constant was achieved with the fluorophore tagged SelK peptide (25 nM, Figure 1B) relative to the reported literature value of 3.4 nM for the unlabeled peptide.^9^ We then demonstrated that unlabeled SelK peptide can compete for KLHDC2 binding (IC_50_ = 55 nM) under these conditions, giving us good confidence in recapitulating this known interaction and providing a positive assay control for high throughput screening (Figure 1C). A suitable 384-well FP assay with GST KLHDC2 (25 nM) and the TAMRA tagged SelK peptide (3.1 nM) was achieved (Z’ = 0.81) when exposed to a molecular excess of free SelK peptide (10 µM; Figure 1D).

The FP assay was further miniaturized to 1536-well format (Z’ = 0.61, Figure 1E) and then screened against subset of the Calibr small molecule library (354,274 compounds) using automated high throughput screening for compounds which decreased the polarization signal (Figure 1F). From ∼14,000 primary screening hits, 51 displayed dose responsive inhibitory signals in follow-up assays. After attriting these molecules to remove potential fluorophores and quenchers, we ultimately identified four validated hit compounds that dose dependently decreased FP signal with half maximal inhibitory concentrations in the single digit micromolar range (Figure 1G).

In parallel, we devised a counter screen to control for nonspecific binders using a different Kelch domain containing ubiquitin E3 ligase, KEAP1 (Kelch-like ECH-associated protein 1), which has also been coopted for targeted protein degradation using PROTACs.^13^ Using recombinant KEAP1 Kelch domain expressed from *E. coli*, we were able to establish a dissociation constant (K_d_ = 22 nM) via FP with a peptide corresponding to the KEAP1 substrate NRF2 (NFE2L2), which was N terminally tagged with a TAMRA fluorophore (TAMRA-AFFAQLQLDEETGEFL, Figure S1A). We confirmed this FP signal could be competed with good potency (IC_50_ = 6 nM) by a previously reported small molecule ligand of the KEAP1 Kelch domain, KI-696 (Figure S1B-C). Importantly, the free SelK peptide displayed no inhibitory signal in FP assays measuring the association between KEAP1 and NRF2 (Figure 1C), further validating the specificity of the SelK-KLHDC2 FP-based interaction assay used for screening.

We then characterized the 4 hits for their capacity to bind to KLHDC2 in vitro. Two of these hits, **1** and **2**, share a conserved 8-carboxy tetahydroquinoline scaffold (Figure 2A, B), whereas the other two compounds, **3** and **4** were closely related imidazopyridines that varied only by an additional methyl substitution present in **4** (Figure S2A, B). The four compounds displayed dose dependent inhibitory effects in the SelK-KLHDC2 competitive FP assay in 384-well format (IC_50_s = 2.5 µM for **1**, 1.3 µM for **2**, 3.6 µM for **3**, and 2.8 µM for **4**) and did not have an inhibitory effect in the KEAP1 counter screening assay at concentrations less than 20 µM (Figure 2C, D; Figure S2C, D). We next evaluated these four molecules for direct binding to immobilized KLHDC2 using surface plasmon resonance (SPR). **1** and **2** gave robust interaction responses, yielding K_d_ values of 810 nM and 440 nM, respectively (Figure 2E, F). Compounds **3** and **4** were not found to provide robust sensorgram signals. Whether this observation derives from incompatibility under the assay conditions or requires the presence of the SelK peptide for interaction with KLHDC2 is unclear. As such, we chose to evaluate the tetrahydroquinoline series for further study.

**Figure 2.**
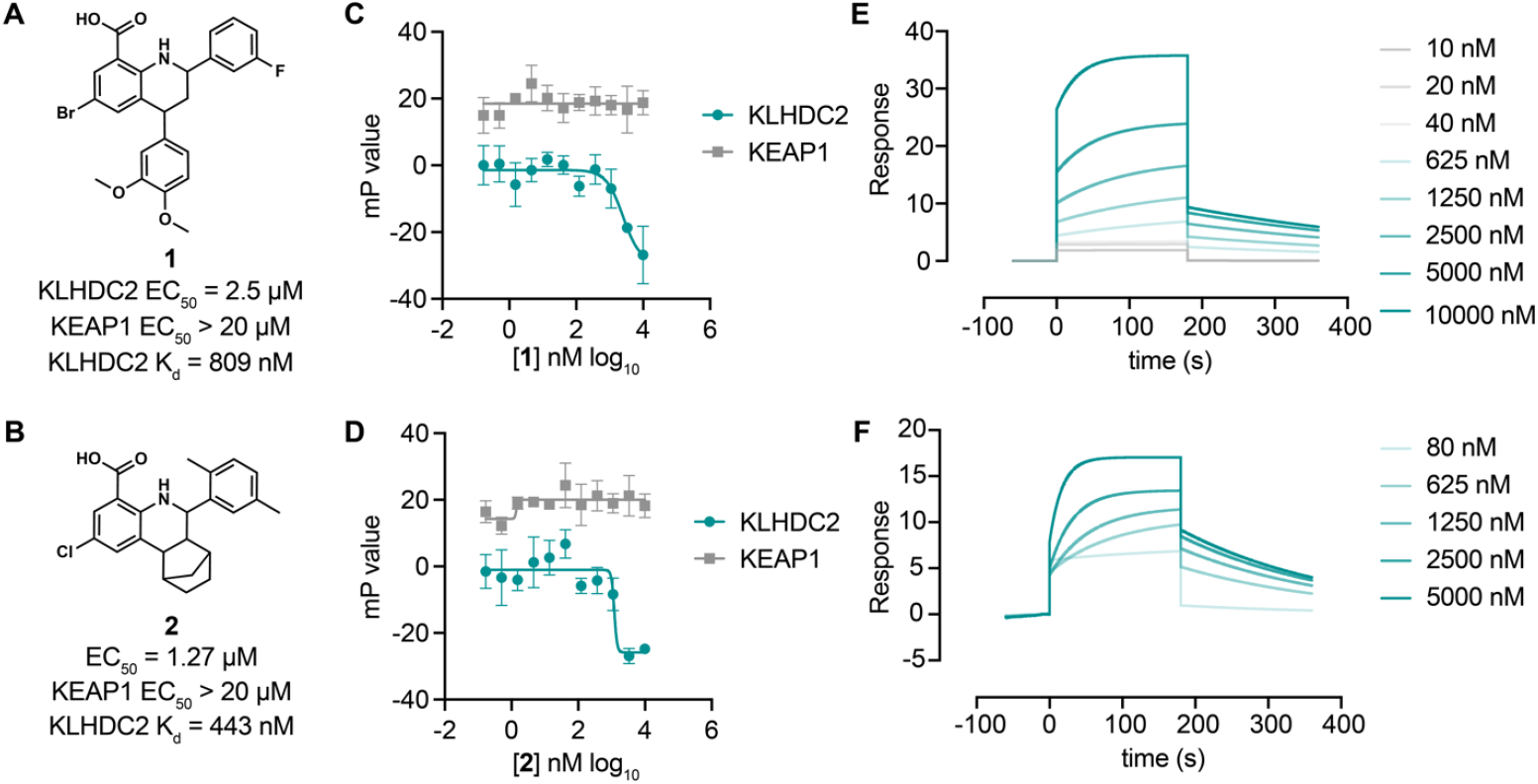
Tetrahydroquinoline-based KLHDC2 ligands. Structures and summary of activities of compounds (A) and **2** (B). C) Fluorescence polarization signal from KLHDC2 and KEAP1 assays in response to increasing concentrations of **1** (C) and **2** (D). Representative SPR sensorgrams of KLHDC2 binding from the indicated concentrations of **1** (E) and **2** (F).

We next sought to understand if the tetrahydroquinoline scaffold might be further optimized for increased affinity with KLHDC2. We evaluated all commercially available analogs that shared the 8-carboxy tetrahydroquinoline core (53 analogs) in 384-well FP assays measuring association between KLHDC2 and the labeled SelK peptide. Notably, substitution of 2,5-dimethyl substituted phenyl group present in **2** to 3-substituted trifluoromethyl or trifluoromethoxy groups was found to increase potency in this initial screen (Figure 3A). We evaluated the trifluoromethyl substituted analog **5** and the trifluoromethoxy substituted **6** (Figure 3B,C) in follow-up FP assays with KLHDC2, observing increased potency for these analogs (IC_50_s = 760 nM and 670 nM respectively, Figure 3D, E) with no inhibitory in the KEAP1 counter screening assay. Compounds **5** and **6** also displayed increased potency in SPR assays measuring affinity for KLHDC2 binding with K_d_s of 380 and 160 nM, respectively (Figure 3F, G).

**Figure 3.**
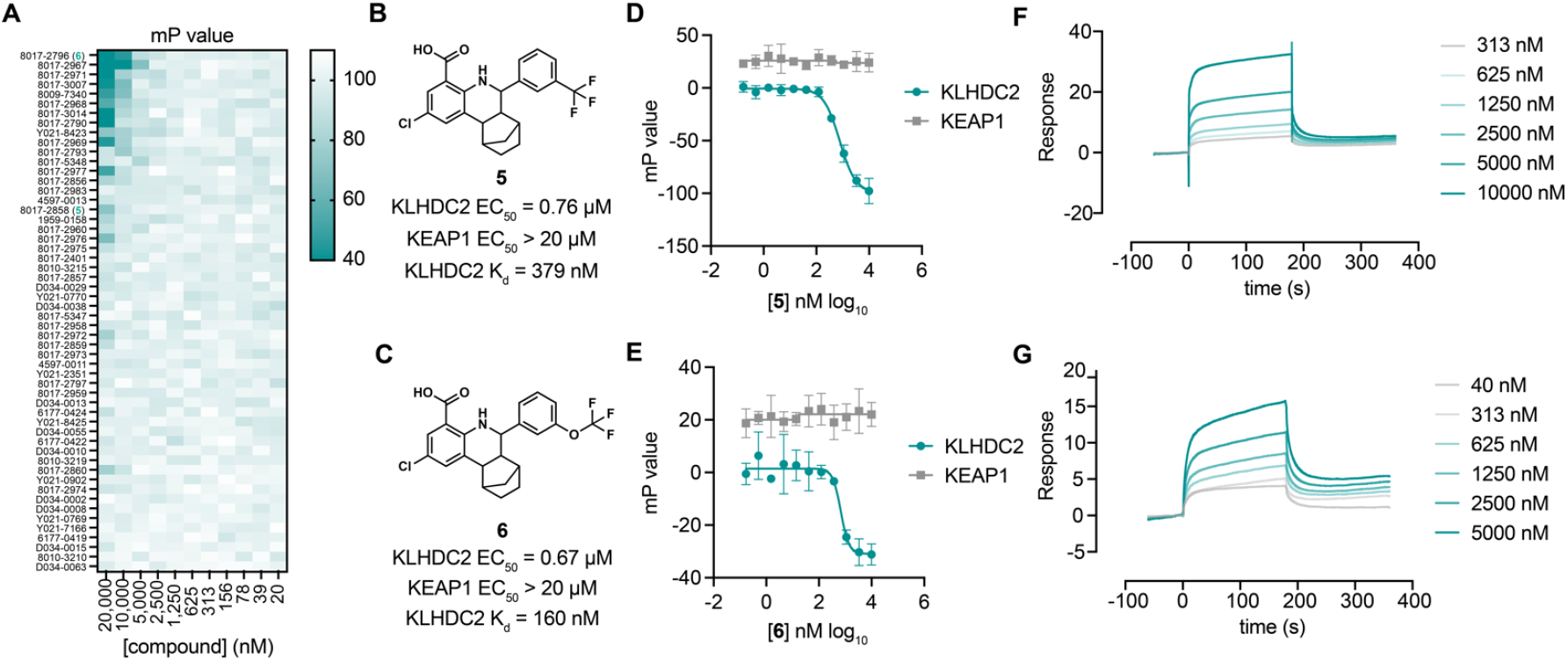
Optimization of tetrahydroquinolines as KLHDC2 ligands. A) Heatmap of fluorescence polarization signal in response to dose responses of the 54 related compounds. Structures and summary of activities of compounds **5** (B) and **6** (C). C) Fluorescence polarization signal from KLHDC2 and KEAP1 assays in response to increasing concentrations of **5** (D) and **6** (E). Representative SPR sensorgrams of KLHDC2 binding from the indicated concentrations of **5** (F) and **6** (G).

We next sought to determine if the tetrahydroquinoline series could be used for targeted protein degradation. We performed docking studies using Autodock Vina to evaluate potential binding modes of **6** to the previously solved crystal structure of the Kelch domain of KLHDC2 (PDB 6DO3).^9^ Surprisingly, **6** was not predicted to be capable of making interactions with Arg236 or Arg241, which recognize the anionic carboxy terminus of the C-terminal diglycine motif of SelK, as anticipated. Instead, **6** occupies a more distal, lipophilic spot in the SelK binding pocket, making a predicted H-bond interaction with Arg189 of KLHDC2 (Figure 4A). In this orientation, the trifluoromethoxy group points to solvent, providing a potential vector for derivatizing PROTAC linkers. We evaluated this hypothesis by synthesizing three PROTAC molecules bearing JQ1, a well characterized ligand to BRD4 (Bromodomain-containing protein 4), which has been used often in the literature as a means of testing PROTAC activity.^6,14^ These molecules were based on the methyl ester substituted scaffold represented by **2** (for increased cellular permeability and subsequent hydrolysis the carboxylate by cellular esterases) and bore 3 position substituted PROTAC linkages to JQ1. These PROTACs linkers consisted of the flexible three ethylene glycol substituted **7** as well as the more structured linkers represented by **8** and **9** (Figure 4B). We evaluated the ability of these molecules to degrade BRD4 using a HEK293T-based assay in which the luminescence produced from a transiently transfected BRD4-HiBiT transgene correlates to the amount of cellular BRD4 present. We found **7** degraded BRD4 in the BRD4-HiBiT with modest activity after 4 hours of treatment (half maximal degradation concentration, DC_50_ = 2.6 µM) and that degradation by this molecule persisted for 24 hours (DC_50_ = 4 µM; Figure 4C, D). Notably, the more rigid PROTAC **8** displayed considerably enhanced activity in degrading BRD4 within 4 hours of treatment (DC_50_ = 164 nM) that further increased (DC_50_ = 80 nM) at the 24-hour timepoint (Figure 4C, D). This result contrasts with dBET1, a reported CRBN-targeting PROTAC degrader of BRD4,^6^ which potently promoted degradation at 4 hours in this assay (DC_50_ = 22 nM) but became inactive at the 24-hour timepoint. Notably, the most rigid biphenyl linker containing **9** did not promote BRD4 degradation at any concentration (< 20 µM) or timepoint (4 or 24 hours) tested. We lastly confirmed the ability of **8** to degrade endogenous BRD4 in SK-BR-3 cells. **8** (100 nM) degraded more than 60% of endogenous BRD4 protein in 24 hours of treatment; maximal degradation was observed with the highest concentration tested (10 µM; 94% degradation; Figure 4E). Importantly, the 10 µM condition could be partially rescued when co-treated with MG-132 (29% remaining vs. 4% in the DMSO condition), indicating degradation occurs through a proteasome-mediated mechanism.

**Figure 4.**
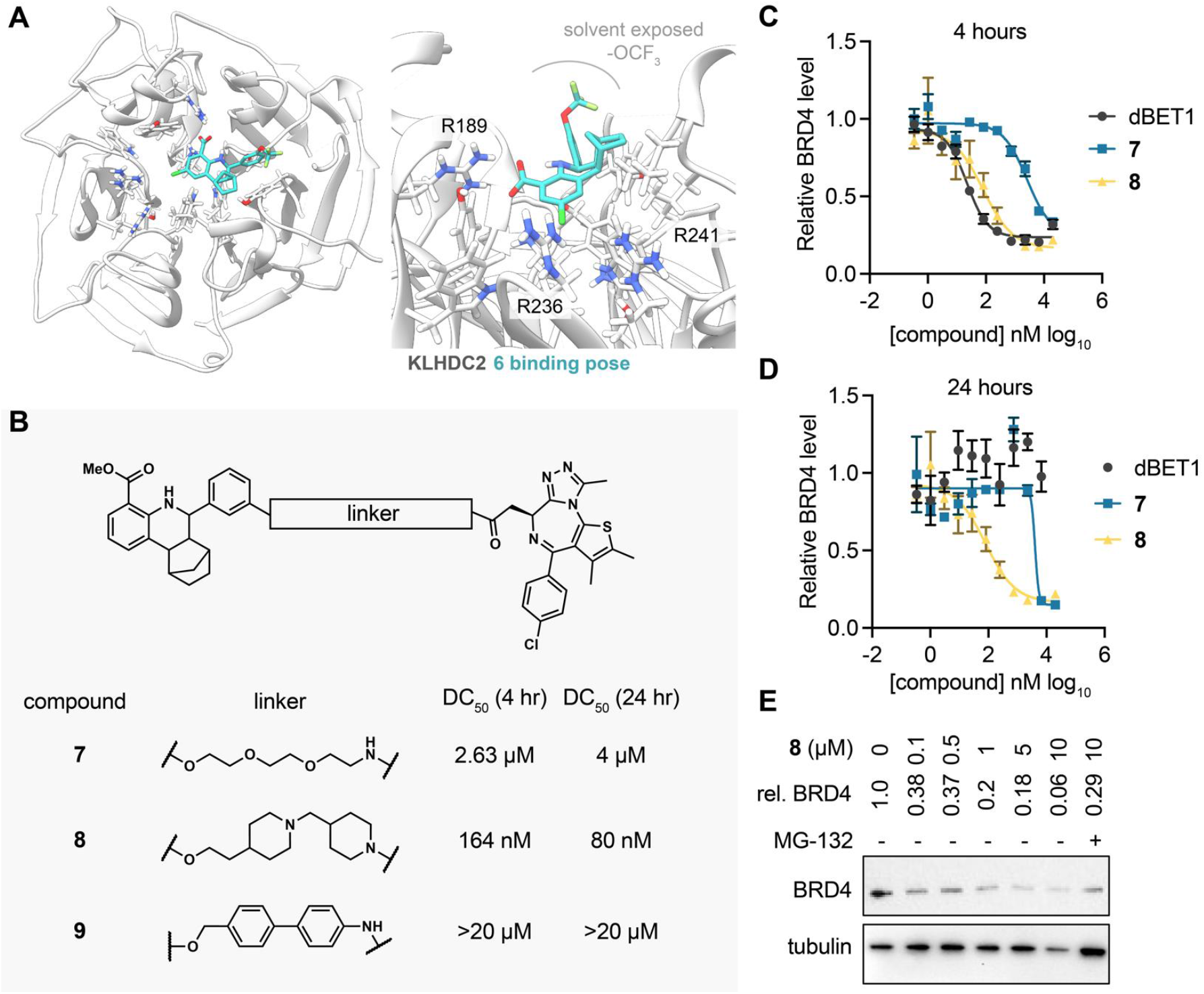
KLHDC2 ligands can be coopted for targeted degradation of BRD4. A) Binding pose of compound 6 to KHLDC2 (PDB 6DO3). B) Structure and summary of activities of the indicated PROTAC molecules. Relative BRD4 levels at 4 (C) or 24 hours (D) as measured by luminescence from HEK293T cells expressing a HiBiT-tagged BRD4 transgene (n=3; mean and s.e.m.). E) Representative Western blot and densitometry-based quantification of BRD4 levels from SK-BR-3 in response to 24-hour treatment with the indicated concentrations of **8**.

Here, we have reported the development and execution of a high-throughput competitive FP-based screen that identified a novel ligand to KLHDC2, a E3 ligase effector of the C-end degradation rule. One tetrahydroquinoline scaffold was found to dose dependently bind KLHDC2 in two in vitro assays, consistent with a binding mode that competes for interaction with KLHDC2 client proteins in the SelK recognition pocket. Importantly, we were able to further optimize the affinity of this series by several fold, yielding **6**, a ligand which bound recombinant KLHDC2 in SPR assays with a K_d_ of 160 nM. As such, we hypothesize that **6** and its analogs will be useful tools to further probe the biological roles of KLHDC2 in cells. Recently, several other groups have used computational design to identify ligands to KLHDC2.^15-17^ These ligands engage the diglycine recognition motif deep within the SelK binding pocket of KLHDC2, which contrasts with the predicted binding mode of **6** presented here. We demonstrated that the tetrahydroquinline scaffold can be further derivatized, yielding PROTAC molecules like **8**, which can degrade BRD4 in cells with long acting degradation kinetics. Like other PROTAC designs, linker geometry is important, with the semi-rigid linker of **8** considerably outperforming other designs. To the extent that **8** may be further modified to increase its potency and drug likeness will be the subject of future work. To our knowledge, this is the first demonstration that a noncovalent, PROTAC-compatible E3 ligase ligand can be discovered by affinity-based, high-throughput screening. Overall, this work adds a novel scaffold for targeting KLHDC2 and further supports the notion that drug like molecules can be developed to this E3 for therapeutic applications.

## Author Contributions

H.Z., T.Z., W.Y., L.L. and Y.K. performed research. K.A.J., I.A.W., M.J.B. oversaw research. H.Z., M.J.B., and P.G.S. wrote the manuscript.

### Notes

The authors declare no competing financial interests.

## Acknowledgements

This work was supported by the NIH (GM146865 to M.J.B. and GM145323 to P.G.S.) and Calibr. We would like to thank Savni Prabhu and Phillip Ordoukhanian for generating SPR data.

## Materials and Methods

### Protein expression and purification

The Kelch domain of human KLHDC2 (amino acids 1-362) was cloned in frame with an N-terminal glutathione-S-transferase (GST) tag and TEV-cleavage sequence into the pFastBac vector and was expressed in two rounds of baculoviral production in insect Sf9 monolayer cells. The produced 15 mL of P3 virus was used to infect 1.5 L HiFive suspension cells at a cellular density of 10^6^/mL for in a 27°C shaker at 110 RPM. After 3 days of infection, the cells were harvested at 8000 x*g* for 30 min and then lysed in 20 mM Tris, pH 8.0, 200 mM NaCl, 5 mM DTT in the presence of protease inhibitors (1 mg/mL Leupeptin, 1 mg/mL Pepstatin and 100 mM PMSF) with sonication. The lysate was incubated with Pierce Glutathione Agarose (Thermo Scientific) at 4°C overnight and eluted with 250mM Glutathione. The crude protein was first dialyzed into dialysis buffer (20 mM Tris, pH 8.0, 200 mM NaCl, 5 mM DTT) and further purified with size-exclusion using a Superdex-75 column (GE Healthcare).The human KEAP1 Kelch domain was expressed using a pET21a-KEAP1 plasmid outfitted cloned in frame with an N-terminal HIS_6_ tag. Plasmid containing BL21(DE3) cells were amplified in 1L 2YT culture at 37 °C until the OD_600_ reached 0.8 followed by 1 mM IPTG induction at 4°C overnight. The harvested cells were lysed in the aforementioned lysis buffer and purified via a standard Ni-NTA purification protocol. Both proteins were concentrated with an Amicon concentrator (Sigma) and the protein concentrations were determined by BCA assay (Thermo fisher 23327). All protein samples were flash frozen in liquid nitrogen for future use.

### Small molecule libraries

The small molecule library of 354,274 compounds consisted of a ChemDiv library (150,114 compounds), an Enamine library (142,208 compounds), a Life Chemicals library (33,792 compounds) and the ReFrame library (28,160 compounds).

### High Throughput Screening

Compounds from selected small molecule libraries were pre-spotted at 10 µM (final concentration) with Echo Acoustic liquid handler system (Beckman) in the format of 1536-well or 384-well in Greiner solid black microplates (Cat.No.782076 and 781209). The C-terminal peptide of SelK, native substrate KLHDC2 protein, HLRGSPPPMAGG (InnoPep. Inc) was spotted into each individual plate which serves as positive control at a final concentration of 1 μM. The FP assay was carried out by first dispensing 5 μL of KLHDC2 protein at 25 nM for 1536-well microplates or 25 μL at 25 nM for 384-well microplates with a Multidrop Dispenser (Themo fisher) followed by incubation at room temperature for 1 h. An equal volume of 3.12 nM TAMRA-HLRGSPPPMAGG (InnoPep. Inc) was dispensed into the plates and incubated at room temperature for another 1 h in the dark. The final concentration of the protein and the corresponding TAMRA-peptide was 12.5 nM and 1.56 nM, respectively. FP signals was recorded using an Envision plate reader (Perkin Elmer) and the robust Z-scores were calculated using Genedata screener where ∼3% of the hits were selected each batch based on robust Z-scores for the primary screening. The second round of triplicate validation and the third round of dose-response characterization were carried out in a similar format.

### Fluorescence polarization assays

The in-house validation of KLHDC binders including re-purchased SAR by inventory compounds was carried out in black solid-bottom 384-well microplates. 25 μL of KLHDC2 or KEAP1 protein was first dispensed into the plates followed by transferring corresponding concentration of compound through Bravo Automated liquid handling system (Aligent). The mixture was incubated at room temperature for 1hr before the TAMRA-peptide was added. The final concentration of the protein and the corresponding TAMRA-peptide was 25 nM and 3.12 nM respectively. The plates were incubated at room temperature in a shaded environment for one more hour and the FP signals were measured by Envision plate reader (Perkin Elmer). The data were analyzed and visualized by GraphPad Prism software.

### SPR binding experiments with KLHDC2^**Klech**^

10 μM His-GST-KLHDC2 in 100 μL of 1X HBS (Hepes buffered saline, 10 mM HEPES, 150 mM NaCl, pH 7.4) containing 25 μM EZ-Link NHS-PEG4-biotin was incubated at 4°C for 2 hours, after which the reaction was quenched with 2 μL of 1 M Tris pH 7.5. The solution was dialyzed over an 18-hour period against 3 changes of 500 mL of 1X HBSS buffer. Biotinylated-His-GST-KLHDC2 was immobilized onto Xantec High Density SA chip by diluting to 0.7 μM injected over the surface for 60 sec at 10 μL/min and resulting in approximate immobilized signal gain of 4,000-4,500 response units (RU). The running buffer for all immobilizations assays was 50 mM Tris pH8, 250 mM NaCl, 0.05% Tween 20, 0.1 mM DTT, and 2% DMSO. All measurements of direct binding in SPR experiments were collected using the Biacore 8K+ instrumentation.

### Cell-based HiBIT BRD4 degradation assay

HET293T cells (0.2 × 10^6^ cells/well, 40 μL) were seeded in white opaque bottom Greiner 384-well microplate for 24 hours before transfection with 1 ng pCMV6-HiBIT-BRD4 plasmid, 99 ng pUC19 plasmid, and 4 µL per µg FuGene (Promega) in OptiMEM (Thermo Fisher). The next day, compounds were spotted as DMSO solutions (100 nL) using a Bravo Automated Liquid Handler (Agilent) affixed with a pintool head (V&P Scientific) in serial dilution. After 4 hours of incubation, 30 μL of Nano-Glo^®^ HiBiT Lytic Reagent was added, and the plates were shaken for 5 min. The HiBIT-BRD4 levels of each well were measured by Envision plate reader (Perkin Elmer). The raw data was analyzed and visualized by GraphPad Prism software.

### Cell-based endogenous BRD4 degradation assay

SK-BR-3 cells (0.25 × 10^6^ cells/well, 2mL) were seeded in 6-well polystyrene plates for 24 hours before the compounds were added at the indicated doses as DMSO solutions. The protease inhibitor MG132 (Selleck chemicals) was added to sixth well along with 10μM 3d at concentration of 10μM. The plate was incubated at 37°C for 24 hours and the cells from each well were collected and lysed in RIPA buffer (Thermo fisher 89900) with Halt protease and phosphatase inhibitor cocktail (Thermo fisher 78440). The cellular content was extracted on ice for 20 min and the protein concentration determined by BCA assay (Thermo fisher 23327). 2 μg of each sample resolved via SDS-PAGE gels and Western blotting used to visualize the endogenous BRD4 level (BRD4 primary antibody Abcam plc, Ab128874, 1:10,000 dilution) with Bio-Rad Gel Doc XR+ imaging system.

## Supporting Information

**Figure S1.**
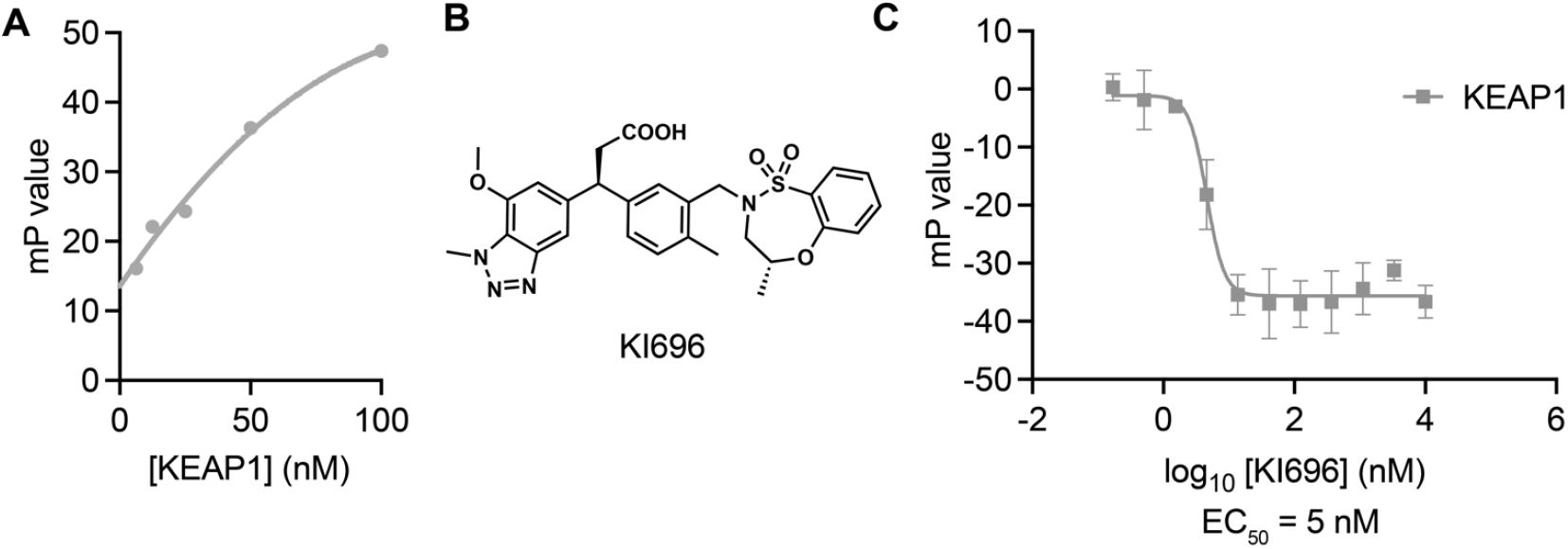
A fluorescence polarization-based control assay with Kelch domain containing protein KEAP1. A) Fluorescence polarization signal of TAMRA-conjugated NRF2 peptide in response to increasing concentrations of KEAP1. B) Structure of KEAP1 Kelch domain ligand KI696. C) Fluorescence polarization signal in response to the indicated concentrations of KI696 in the KEAP1 assay. N=3, mean and s.e.m.

**Figure S2.**
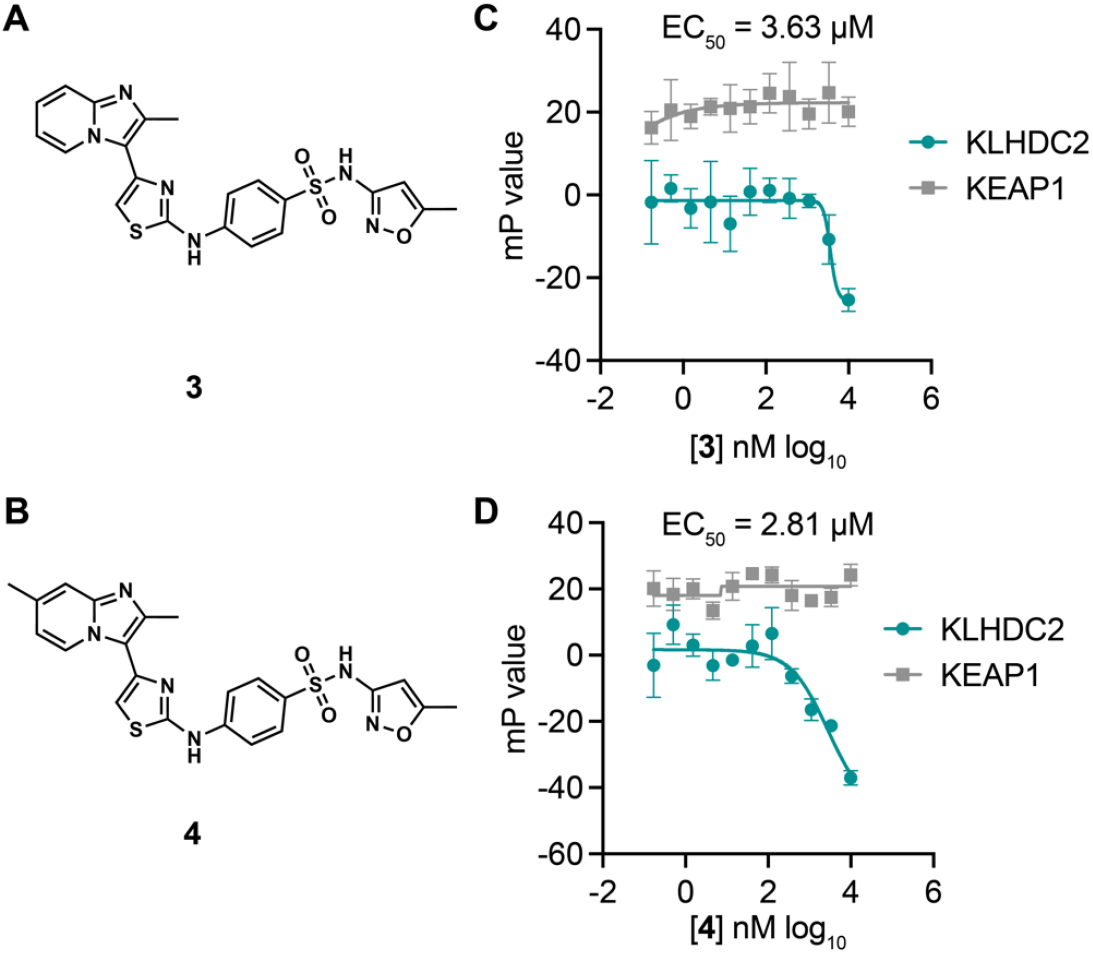
A 2-amino thiazole-based KLHDC2 ligand scaffold. Structures of compounds **3** (A) and **4** (B). Fluorescence polarization signal from KLHDC2 and KEAP1 assays from the indicated concentrations of 3 (C) and 4 (D) (n=3; mean and s.e.m.).

### Synthetic methods and characterization

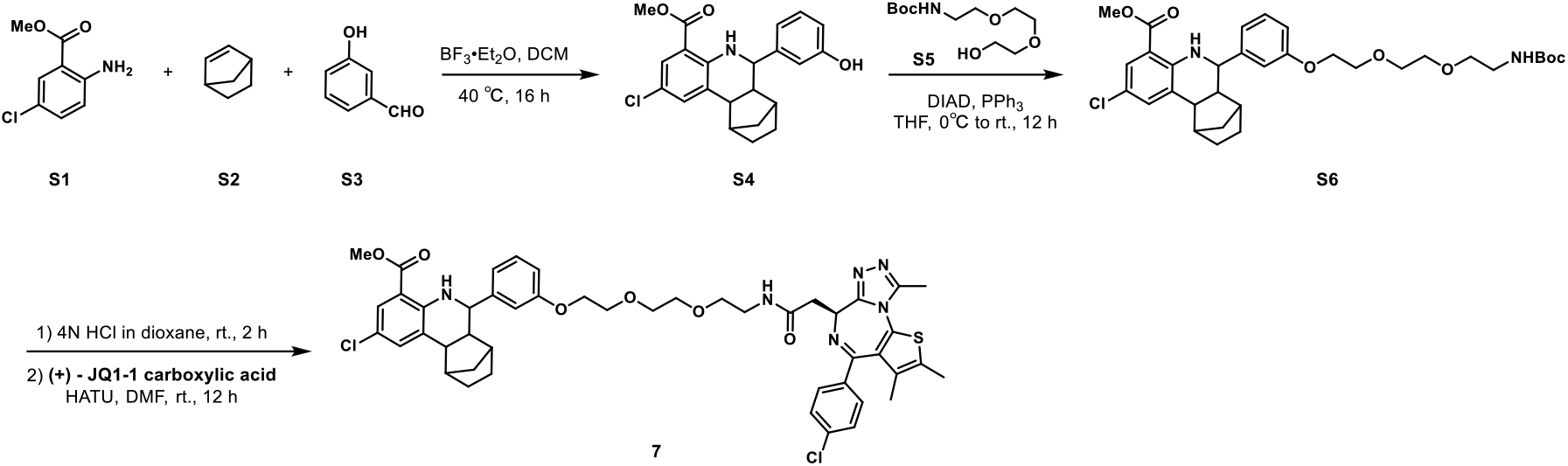

#### Methyl 2-chloro-6-(3-hydroxyphenyl)-5,6,6a,7,8,9,10,10a-octahydro-7,10-methanophenanthridine-4-carboxylate (S4)

Methyl 2-amino-5-chlorobenzoate S1 (1.85 g, 10 mmol), norbornene S2 (1.41 g, 15 mmol) and 3-hydroxybenzaldehyde S3 (1.34 g, 11 mmol) were dissolved in DCM (50 mL) in a 200 mL Ace pressure tube. After purging with argon, boron trifluoride etherate (284 mg, 2 mmol) was added. The tube was sealed instantly, and the reaction mixture was heated at 40 °C for 16 h. 1 mL of methanol was added to the mixture to quench the reaction. After concentrated in vacuo, the crude residue was purified by flash chromatography on silica gel to give S4 (1.75 g, 42% yield) as yellow solid. ^1^H NMR (400 MHz, CDCl_3_) δ 7.72 (s, 1H), 7.50 (s, 1H), 7.38 (s, 1H), 7.21 (t, *J* = 7.7 Hz, 1H), 6.94 (d, *J* = 7.5 Hz, 1H), 6.89 (s, 1H), 6.77 (d, *J* = 7.9 Hz, 1H), 5.09 (s, 1H), 3.80 (s, 3H), 3.56 (d, *J* = 9.7 Hz, 1H), 2.70 – 2.57 (m, 2H), 2.22 – 2.10 (m, 2H), 1.71 – 1.60 (m, 3H), 1.58 – 1.50 (m, 1H), 1.44 – 1.36 (m, 1H), 1.12 (d, *J* = 9.9 Hz, 1H).

#### Methyl 2-chloro-6-(3-((2,2-dimethyl-4-oxo-3,8,11-trioxa-5-azatridecan-13-yl)oxy)phenyl)-5,6,6a,7,8,9,10,10a-octahydro-7,10-methanophenanthridine-4-carboxylate (S6)

S4 (500 mg, 1.3 mmol), tert-butyl (2-(2-(2-hydroxyethoxy)ethoxy)ethyl)carbamate S5 (389 mg, 1.56 mmol) and triphenylphosphine (511 mg, 1.95 mmol) were dissolved in THF (20 mL) in a 100 mL flask. The reaction mixture was cooled to 0 °C with ice water bath. Diisopropyl azodicarboxylate (DIAD, 394 mg, 1.95 mmol) was added dropwise, after which the reaction mixture was allowed to warm to room temperature and stirred for 16 hours. The mixture was quenched with aqueous NH_4_Cl solution (50 mL). The aqueous layer was extracted with ethyl acetate (100 mL × 2), and the combined organic layers were washed with brine (100 mL × 2), dried over anhydrous Na_2_SO_4_, filtered and concentrated in vacuo. The crude residue was purified by flash chromatography on silica gel to give S6 (688 mg, 86% yield) as yellow solid. ^1^H NMR (400 MHz, CDCl_3_) δ 7.72 (dd, *J* = 2.5, 0.8 Hz, 1H), 7.50 (s, 1H), 7.38 (s, 1H), 7.26 – 7.21 (m, 1H), 7.01 – 6.95 (m, 2H), 6.86 (dd, *J* = 8.2, 1.6 Hz, 1H), 5.03 (s, 1H), 4.18 – 4.14 (m, 2H), 3.89 – 3.85 (m, 2H), 3.80 (s, 3H), 3.75 – 3.70 (m, 2H), 3.67 – 3.63 (m, 2H), 3.58 – 3.53 (m, 3H), 3.35 – 3.29 (m, 2H), 2.67 – 2.60 (m, 2H), 2.20 – 2.12 (m, 2H), 1.65 – 1.60 (m, 4H), 1.44 – 1.41 (m, 10H), 1.13 (d, *J* = 10.2 Hz, 1H).

#### Methyl 2-chloro-6-(3-(2-(2-(2-(2-((*S*)-4-(4-chlorophenyl)-2,3,9-trimethyl-6*H*-thieno[3,2-*f*][1,2,4]triazolo[4,3-*a*][1,4]diazepin-6-yl)acetamido)ethoxy)ethoxy)ethoxy)phenyl)-5,6,6a,7,8,9,10,10a-octahydro-7,10-methanophenanthridine-4-carboxylate (7)

S6 (92 mg, 0.15 mmol) was dissolved in 4 M HCl in dioxane (5 mL) which was then stirred at room temperature for 2 hours. The mixture was then concentrated under reduced pressure. To the solution of the above residue and (+)-JQ1 carboxylic acid (40 mg, 0.1 mmol) in DMF (5 mL) was added diisopropylethylamine (77.4 mg, 0.6 mmol) and HATU (57 mg, 0.15 mmol). The mixture was stirred at 25 °C for 12 h, then diluted with 1 M aqueous HCl (20 mL) and extracted with ethyl acetate (20 mL × 2). The combined organic layers were washed with aqueous NaHCO_3_ solution, and brine (20 mL × 2), dried over sodium sulfate, filtered, and concentrated. The crude residue was purified by preparative HPLC to give 7 (19 mg, 21% yield) as yellow solid. ^1^H NMR (400 MHz, CDCl_3_) δ 7.71 (d, *J* = 2.3 Hz, 1H), 7.44 – 7.32 (m, 6H), 7.22 (dd, *J* = 7.9, 1.9 Hz, 1H), 6.97 (d, *J* = 7.5 Hz, 2H), 6.86 (d, *J* = 8.0 Hz, 1H), 4.78 (t, *J* = 7.0 Hz, 1H), 4.17 (t, *J* = 4.6 Hz, 2H), 3.90 (d, *J* = 2.9 Hz, 2H), 3.79 (s, 3H), 3.78 – 3.74 (m, 2H), 3.73 – 3.69 (m, 2H), 3.68 – 3.60 (m, 2H), 3.59 – 3.49 (m, 4H), 3.48 – 3.42 (m, 1H), 2.82 (s, 3H), 2.65 (d, *J* = 8.9 Hz, 1H), 2.61 (d, *J* = 3.4 Hz, 1H), 2.44 (s, 3H), 2.19 (t, *J* = 9.1 Hz, 1H), 2.15 – 2.10 (m, 1H), 1.69 (s, 3H), 1.67 – 1.58 (m, 2H), 1.57 – 1.47 (m, 1H), 1.45 – 1.36 (m, 1H), 1.26 – 1.16 (m, 1H), 1.12 (d, J = 10.1 Hz, 1H). ^13^C NMR (101 MHz, CDCl_3_) δ 171.03, 167.97, 159.05, 150.68, 149.16, 145.30, 138.25, 135.10, 133.49, 133.17, 131.85, 131.25, 131.11, 130.53, 130.32, 129.79, 129.77, 129.16, 128.04, 121.50, 120.57, 120.51, 114.00, 113.97, 112.76, 70.89, 70.53, 69.88, 69.53, 67.53, 59.73, 53.66, 52.75, 51.91, 43.96, 43.63, 40.09, 40.01, 37.79, 34.02, 29.77, 29.14, 14.45, 13.35, 11.30. HRMS calcd for C_47_H_51_Cl_2_N_6_O_6_S (M^+^ + H) 897.2968, found 897.3001.

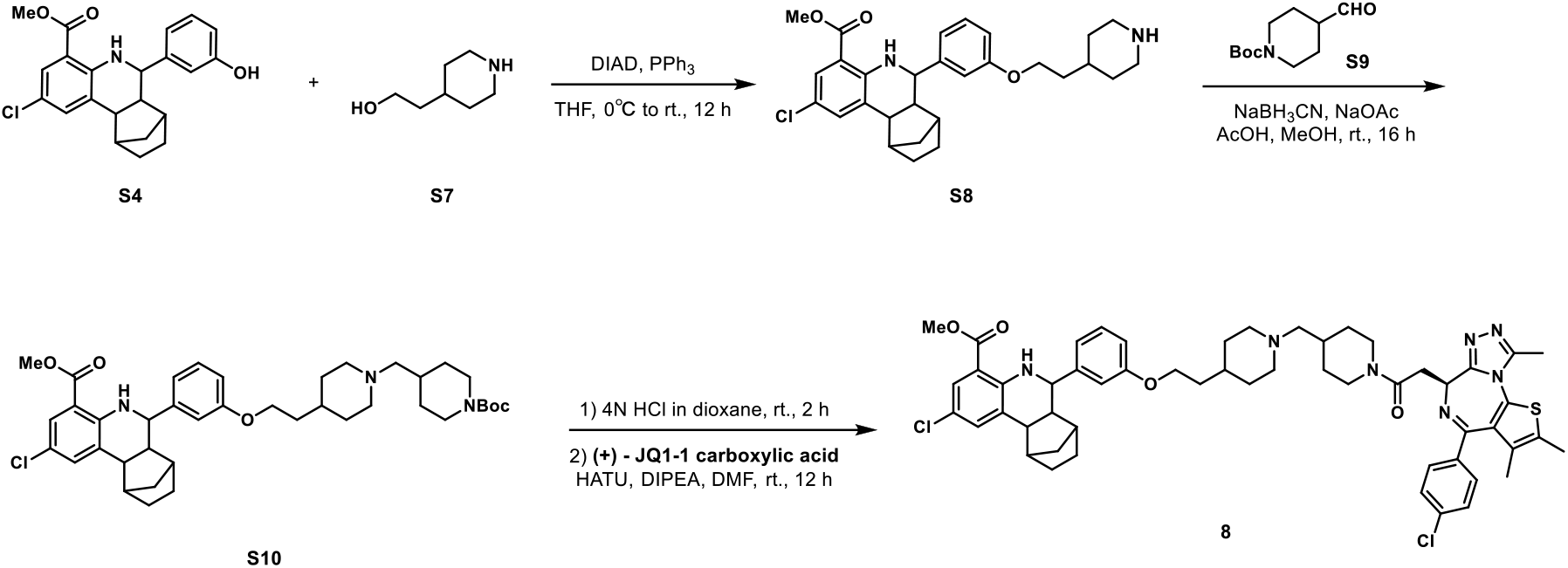

#### Methyl 2-chloro-6-(3-(2-(piperidin-4-yl)ethoxy)phenyl)-5,6,6a,7,8,9,10,10a-octahydro-7,10-methanophenanthridine-4-carboxylate (S8)

S4 (766 mg, 2 mmol), 2-(piperidin-4-yl)ethan-1-ol S7 (307 mg, 2.4 mmol) and triphenylphosphine (786 mg, 3 mmol) were dissolved in THF (25 mL) in a 100 mL flask. The reaction mixture was cooled to 0 °C with ice water bath. Diisopropyl azodicarboxylate (DIAD, 606 mg, 3 mmol) was added dropwise, after which the reaction mixture was allowed to warm to room temperature and stirred for 16 hours. The mixture was quenched with aqueous NH_4_Cl solution (50 mL). The aqueous layer was extracted with ethyl acetate (40 mL × 2), and the combined organic layers were washed with brine (40 mL × 2), dried over anhydrous Na_2_SO_4_, filtered and concentrated in vacuo. The crude residue was purified by flash chromatography on silica gel to give S8 (574 mg, 58% yield) as yellow solid. ^1^H NMR (400 MHz, CDCl_3_) δ 7.75 (dd, *J* = 2.5, 0.8 Hz, 1H), 7.41 (dd, *J* = 2.4, 1.2 Hz, 1H), 7.28 – 7.25 (m, 1H), 7.00 (d, *J* = 7.7 Hz, 1H), 6.99 – 6.96 (m, 1H), 6.84 (dd, *J* = 8.2, 1.9 Hz, 1H), 4.04 (t, *J* = 5.8 Hz, 2H), 3.83 (s, 3H), 3.59 (d, *J* = 9.9 Hz, 1H), 3.45 (d, *J* = 12.0 Hz, 2H), 3.00 – 2.85 (m, 2H), 2.70 – 2.62 (m, 2H), 2.22 (t, *J* = 9.4 Hz, 1H), 2.17 (d, *J* = 3.8 Hz, 1H), 2.00 (d, *J* = 13.9 Hz, 2H), 1.95 – 1.87 (m, 1H), 1.86 – 1.79 (m, 2H), 1.73 – 1.53 (m, 5H), 1.49 – 1.40 (m, 1H), 1.30 – 1.20 (m, 1H), 1.16 (d, *J* = 10.2 Hz, 1H).

#### Methyl 6-(3-(2-(1-((1-(*tert*-butoxycarbonyl)piperidin-4-yl)methyl)piperidin-4-yl)ethoxy)phenyl)-2-chloro-5,6,6a,7,8,9,10,10a-octahydro-7,10-methanophenanthridine-4-carboxylate (S10)

To a solution of S8 (99 mg, 0.2 mmol) in methanol (3 mL) was added sodium acetate (29 mg, 0.4 mmol), acetic acid (10 µL, 0.2 mmol) and sodium cyanoborohydride (33 mg, 0.5 mmol) at 0 °C. Then tert-butyl 4-formylpiperidine-1-carboxylate S9 (56 mg, 0.3 mmol) was added and the mixture and stirred at room temperature for 16 h. The reaction was concentrated under reduced pressure. The residue was purified by flash chromatography on silica gel to give S10 (101 mg, 71% yield) as yellow solid. ^1^H NMR (400 MHz, CDCl_3_) δ 7.72 (d, *J* = 1.7 Hz, 1H), 7.48 (s, 1H), 7.39 (s, 1H), 7.26 – 7.22 (m, 1H), 6.97 (d, *J* = 7.7 Hz, 1H), 6.95 – 6.92 (m, 1H), 6.82 (dd, *J* = 8.2, 1.9 Hz, 1H), 4.20 – 4.06 (m, 2H), 4.01 (t, *J* = 5.8 Hz, 2H), 3.80 (s, 3H), 3.56 (d, *J* = 9.8 Hz, 1H), 3.27 – 3.08 (m, 2H), 2.72 – 2.61 (m, 4H), 2.19 (t, *J* = 9.4 Hz, 1H), 2.14 (d, *J* = 3.8 Hz, 1H), 1.90 – 1.84 (m, 2H), 1.80 – 1.70 (m, 5H), 1.70 – 1.50 (m, 10H), 1.45 (s, 9H), 1.41 – 1.37 (m, 1H), 1.24 – 1.19 (m, 1H), 1.18 – 1.15 (m, 1H), 1.15 – 1.10 (m, 2H).

#### Methyl 2-chloro-6-(3-(2-(1-((1-(2-((*S*)-4-(4-chlorophenyl)-2,3,9-trimethyl-6*H*-thieno[3,2-*f*][1,2,4]triazolo[4,3-*a*][1,4]diazepin-6-yl)acetyl)piperidin-4-yl)methyl)piperidin-4-yl)ethoxy)phenyl)-5,6,6a,7,8,9,10,10a-octahydro-7,10-methanophenanthridine-4-carboxylate (8)

S10 (72 mg, 0.1 mmol) was dissolved in 4 M HCl in dioxane (3 mL) which was then stirred at room temperature for 2 hours. The mixture was then concentrated under reduced pressure. To the solution of the above residue and (+)-JQ1 carboxylic acid (49 mg, 0.12 mmol) in DMF (5 mL) was added diisopropylethylamine (65 mg, 0.5 mmol) and HATU (57 mg, 0.15 mmol). The mixture was stirred at room temperature for 12 h, then diluted with 1 M aqueous HCl (20 mL) and extracted with ethyl acetate (20 mL × 2). The combined organic layers were washed with aqueous NaHCO_3_ solution, and brine (20 mL × 2), dried over sodium sulfate, filtered, and concentrated. The crude residue was purified by preparative HPLC to give 8 (15 mg, 15% yield) as yellow solid. ^1^H NMR (400 MHz, CDCl_3_) δ 7.72 (s, 1H), 7.43 – 7.35 (m, 5H), 7.26 – 7.22 (m, 1H), 6.98 (d, *J* = 7.7 Hz, 1H), 6.94 (s, 1H), 6.83 – 6.80 (m, 1H), 4.91 – 4.83 (m, 1H), 4.64 – 4.53 (m, 1H), 4.29 – 4.13 (m, 1H), 4.06 – 3.98 (m, 2H), 3.97 – 3.82 (m, 1H), 3.80 (s, 3H), 3.78 – 3.68 (m, 2H), 3.57 (d, *J* = 9.9 Hz, 1H), 3.35 – 3.14 (m, 2H), 3.06 – 2.90 (m, 2H), 2.82 (s, 3H), 2.78 – 2.57 (m, 5H), 2.44 (s, 3H), 2.30 – 2.16 (m, 2H), 2.15 – 2.12 (m, 1H), 2.06 – 1.93 (m, 3H), 1.90 – 1.74 (m, 6H), 1.70 (s, 3H), 1.69 – 1.59 (m, 2H), 1.59 – 1.50 (m, 1H), 1.43 – 1.38 (m, 1H), 1.31 – 1.18 (m, 2H), 1.14 (d, *J* = 10.1 Hz, 1H). ^13^C NMR (101 MHz, CDCl_3_) δ 168.63, 168.07, 168.04, 165.37, 164.99, 160.95, 160.56, 159.09, 155.54, 149.33, 145.47, 137.76, 135.73, 133.50, 133.09, 131.78, 131.67, 130.43, 130.34, 130.07, 129.77, 129.14, 129.09, 128.05, 121.40, 120.53, 114.28, 113.72, 113.68, 112.66, 64.60, 63.90, 59.74, 55.09, 54.85, 54.54, 52.86, 51.89, 46.25, 43.97, 43.59, 42.26, 39.99, 34.92, 34.57, 34.03, 32.37, 30.74, 30.70, 30.35, 29.79, 29.15, 29.02, 14.47, 13.35, 11.16. HRMS calcd for C_54_H_62_Cl_2_N_7_O_4_S (M^+^ + H) 974.3961, found 974.3993.

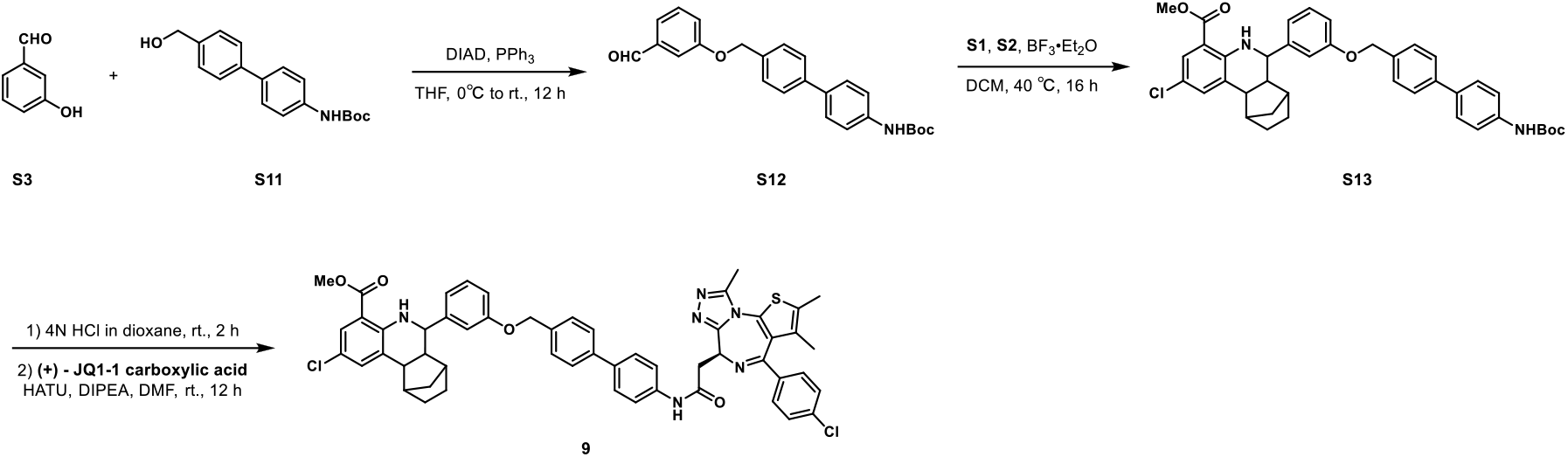

#### *tert*-Butyl (4’-((3-formylphenoxy)methyl)-[1,1’-biphenyl]-4-yl)carbamate (S12)

S3 (244 mg, 2 mmol), tert-butyl (4’-(hydroxymethyl)-[1,1’-biphenyl]-4-yl)carbamate S11 (1.3 g, 2.4 mmol) and triphenylphosphine (629 mg, 2.4 mmol) were dissolved in THF (10 mL) in a 50 mL flask. The reaction mixture was cooled to 0 °C with ice water bath. Diisopropyl azodicarboxylate (DIAD, 485 mg, 2.4 mmol) was added dropwise, after which the reaction mixture was allowed to warm to room temperature and stirred for 16 hours. The mixture was quenched with aqueous NH_4_Cl solution (50 mL). The aqueous layer was extracted with ethyl acetate (40 mL × 2), and the combined organic layers were washed with brine (40 mL × 2), dried over anhydrous Na_2_SO_4_, filtered and concentrated in vacuo. The crude residue was purified by flash chromatography on silica gel to give S12 (460 mg, 57% yield) as white solid. ^1^H NMR (500 MHz, CDCl_3_) δ 9.98 (s, 1H), 7.59 (d, *J* = 8.1 Hz, 2H), 7.53 (d, *J* = 8.5 Hz, 2H), 7.51 – 7.48 (m, 4H), 7.46 (t, *J* = 6.8 Hz, 2H), 7.29 – 7.26 (m, 1H), 6.55 (s, 1H), 5.16 (s, 2H), 1.54 (s, 9H).

#### Methyl 6-(3-((4’-((*tert*-butoxycarbonyl)amino)-[1,1’-biphenyl]-4-yl)methoxy)phenyl)-2-chloro-5,6,6a,7,8,9,10,10a-octahydro-7,10-methanophenanthridine-4-carboxylate (S13)

Methyl 2-amino-5-chlorobenzoate S1 (185 mg, 1 mmol), norbornene S2 (141 mg, 1.5 mmol) and S12 (443 mg, mmol) were dissolved in DCM (5 mL) in a 50 mL Ace pressure tube. After purging with argon, boron trifluoride etherate (28 mg, 0.2 mmol) was added. The tube was sealed instantly, and the reaction mixture was heated at 40 °C for 16 h. 1 mL of methanol was added to the mixture to quench the reaction. After concentrated in vacuo, the crude residue was purified by flash chromatography on silica gel to give S13 (193 mg, 29% yield) as yellow solid. ^1^H NMR (400 MHz, CDCl_3_) δ 7.72 (d, *J* = 1.9 Hz, 1H), 7.58 (d, *J* = 8.2 Hz, 2H), 7.53 (d, *J* = 8.5 Hz, 3H), 7.49 (d, *J* = 8.2 Hz, 2H), 7.43 (d, *J* = 8.5 Hz, 2H), 7.38 (s, 1H), 7.30 – 7.23 (m, 2H), 7.06 (s, 1H), 7.00 (d, *J* = 7.7 Hz, 1H), 6.93 (dd, *J* = 8.2, 2.0 Hz, 1H), 6.55 (s, 1H), 5.10 (s, 2H), 3.80 (s, 3H), 3.59 (d, *J* = 9.8 Hz, 1H), 2.69 – 2.59 (m, 2H), 2.22 – 2.12 (m, 2H), 1.72 – 1.56 (m, 4H), 1.54 (s, 9H), 1.23 – 1.16 (m, 1H), 1.12 (d, *J* = 10.2 Hz, 1H).

#### Methyl 2-chloro-6-(3-((4’-(2-((*S*)-4-(4-chlorophenyl)-2,3,9-trimethyl-6*H*-thieno[3,2-*f*][1,2,4]triazolo[4,3-*a*][1,4]diazepin-6-yl)acetamido)-[1,1’-biphenyl]-4-yl)methoxy)phenyl)-5,6,6a,7,8,9,10,10a-octahydro-7,10-methanophenanthridine-4-carboxylate (9)

S13 (66 mg, 0.1 mmol) was dissolved in 4 M HCl in dioxane (3 mL) which was then stirred at room temperature for 2 hours. The mixture was then concentrated under reduced pressure. To the solution of the above residue and (+)-JQ1 carboxylic acid (49 mg, 0.12 mmol) in DMF (5 mL) was added diisopropylethylamine (65 mg, 0.5 mmol) and HATU (57 mg, 0.15 mmol). The mixture was stirred at room temperature for 12 h, then diluted with 1 M aqueous HCl (20 mL) and extracted with ethyl acetate (20 mL × 2). The combined organic layers were washed with aqueous NaHCO_3_ solution, and brine (20 mL × 2), dried over sodium sulfate, filtered, and concentrated. The crude residue was purified by preparative HPLC to give 9 (36 mg, 38% yield) as yellow solid. ^1^H NMR (400 MHz, CDCl_3_) δ 8.81 (s, 1H), 7.72 (d, *J* = 1.7 Hz, 1H), 7.64 (d, *J* = 8.6 Hz, 2H), 7.60 – 7.55 (m, 4H), 7.50 (d, J = 8.2 Hz, 2H), 7.42 (d, *J* = 8.5 Hz, 2H), 7.39 – 7.36 (m, 3H), 7.29 – 7.26 (m, 2H), 7.06 (s, 1H), 7.00 (d, *J* = 7.6 Hz, 1H), 6.93 (dd, *J* = 8.2, 1.9 Hz, 1H), 5.10 (s, 2H), 4.81 (t, *J* = 6.8 Hz, 1H), 3.80 (s, 3H), 3.61 – 3.58 (m, 3H), 2.80 (s, 3H), 2.68 – 2.60 (m, 2H), 2.45 (s, 3H), 2.20 (t, *J* = 9.3 Hz, 1H), 2.14 – 2.11 (m, 1H), 1.73 – 1.70 (m, 3H), 1.68 – 1.60 (m, 2H), 1.55 – 1.50 (m, 1H), 1.43 – 1.37 (m, 1H), 1.22 – 1.17 (m, 1H), 1.12 (d, *J* = 10.3 Hz, 1H). ^13^C NMR (101 MHz, CDCl_3_) δ 168.66, 167.99, 159.14, 150.57, 149.35, 145.46, 140.51, 140.36, 137.91, 137.21, 137.18, 135.99, 135.64, 133.49, 132.64, 131.63, 131.27, 130.43, 130.24, 130.22, 130.18, 129.79, 129.14, 128.26, 128.09, 128.06, 127.76, 127.20, 121.34, 120.50, 114.26, 114.23, 112.67, 69.92, 59.71, 54.30, 52.85, 51.90, 43.98, 43.69, 40.06, 39.83, 34.04, 29.80, 29.20, 14.57, 13.36, 11.55. HRMS calcd for C_54_H_49_Cl_2_N_6_O_4_S (M^+^ + H) 947.2908, found 947.2921.

^1^H NMR spectra of **7** in CDCl_3_

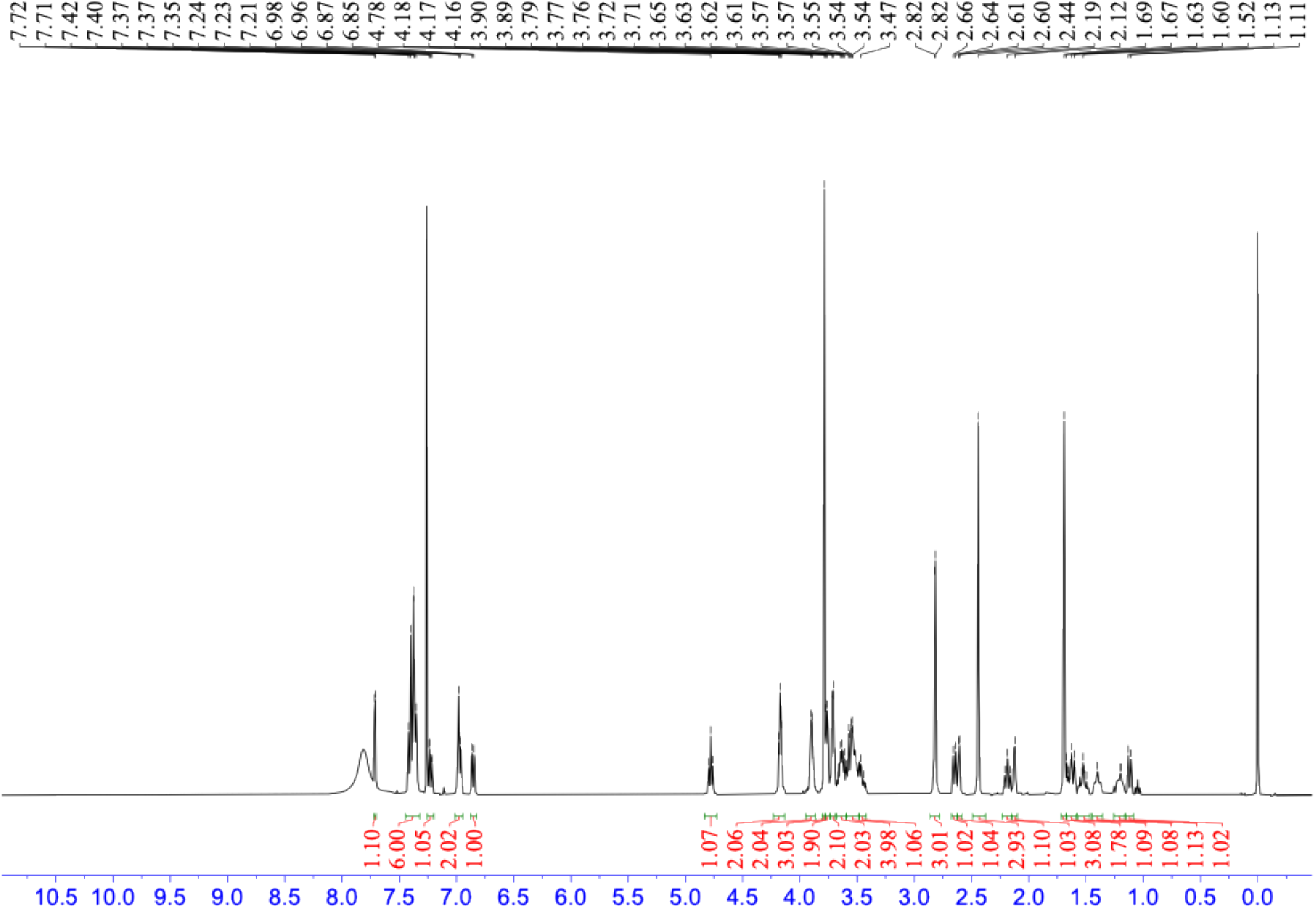

HRMS spectra of **7**

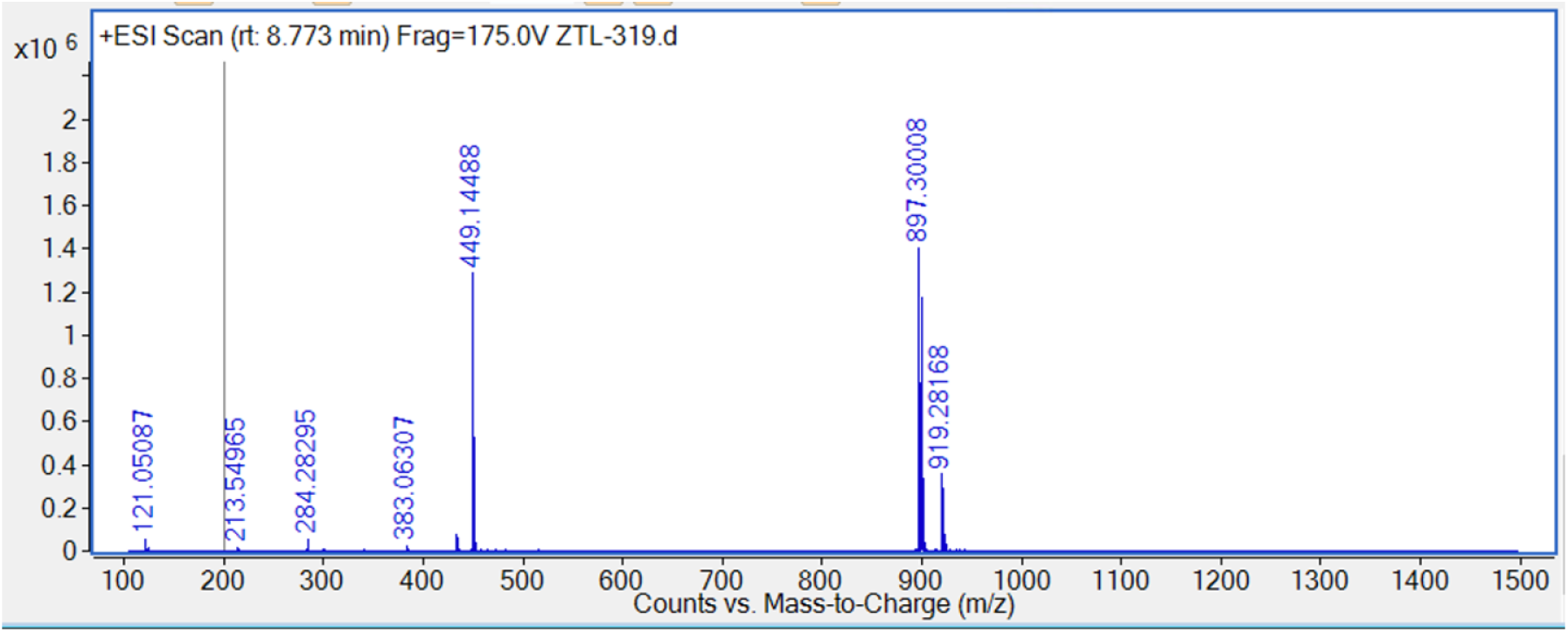

^1^H NMR spectra of **8** in CDCl_3_

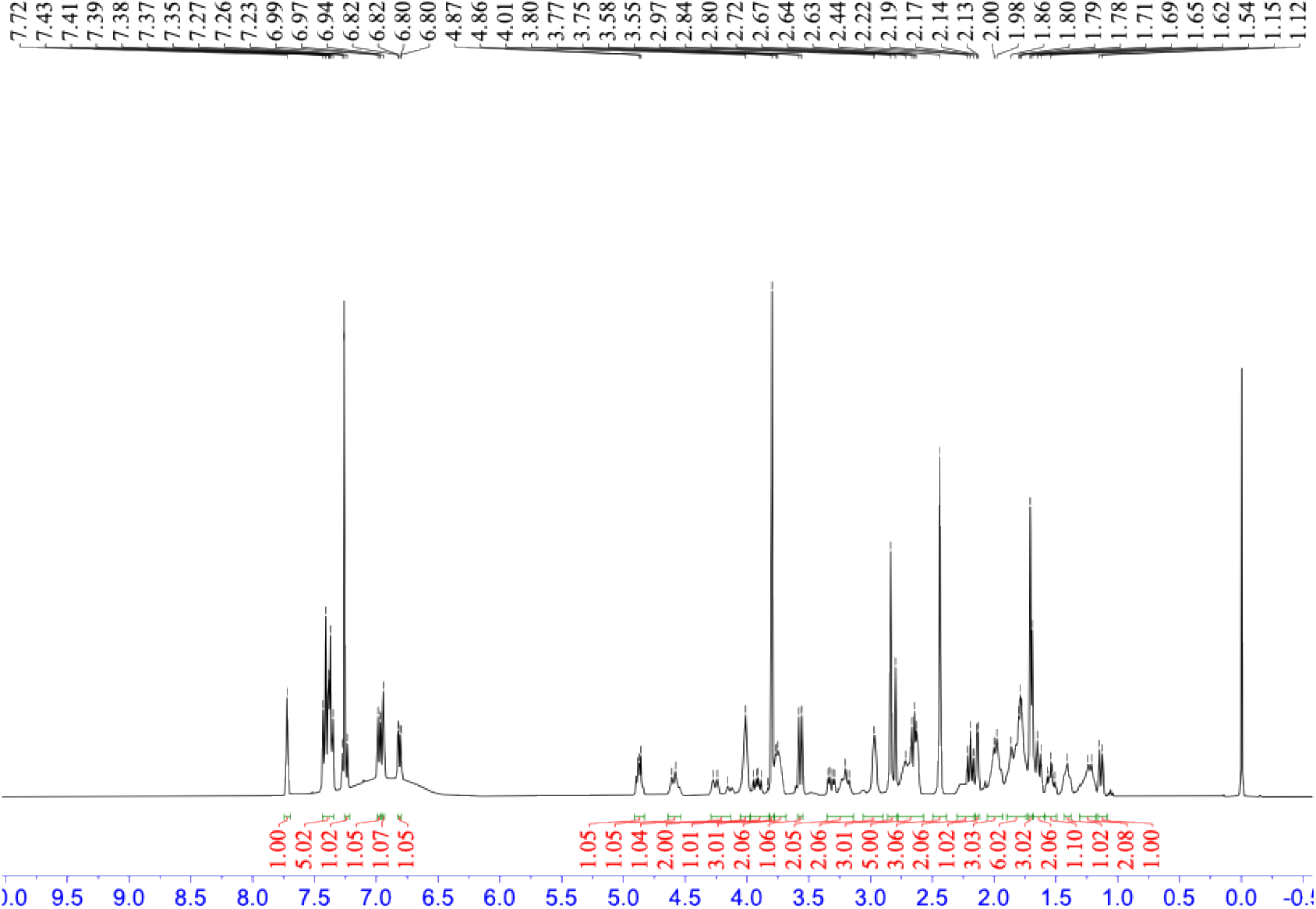

HRMS spectra of **8**

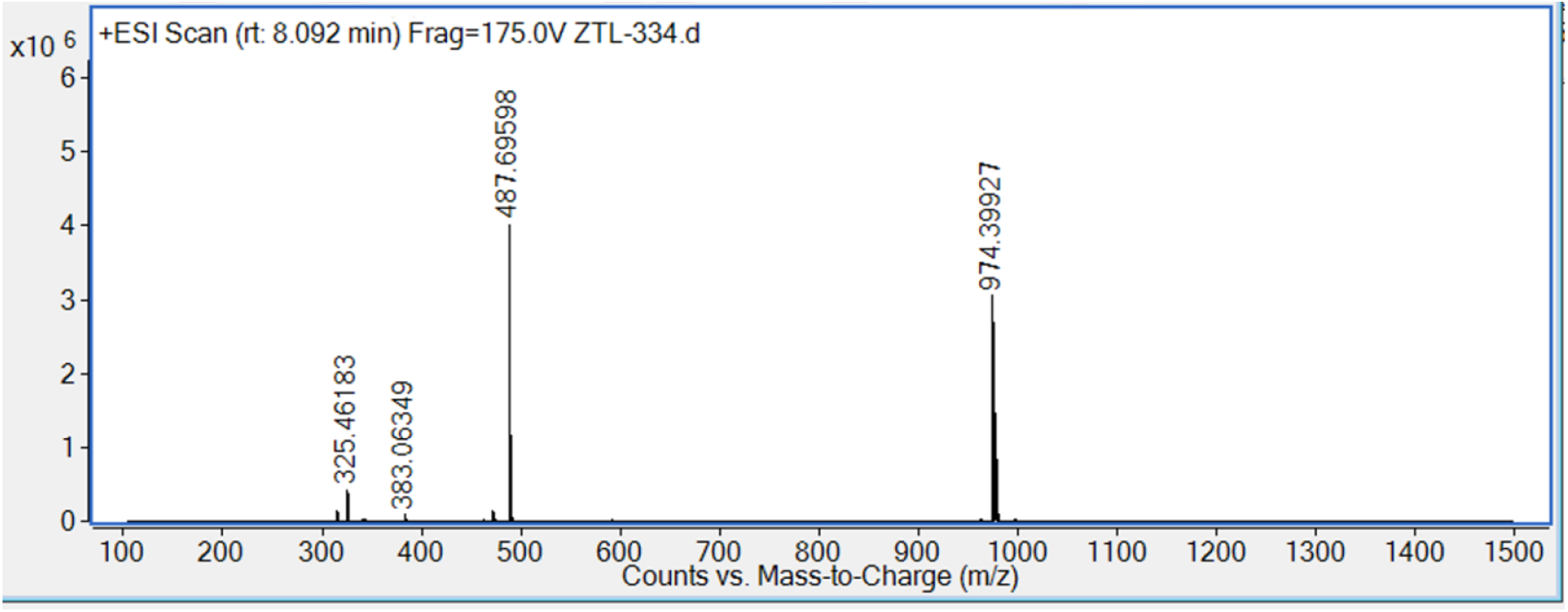

^1^H NMR spectra of **9** in CDCl_3_

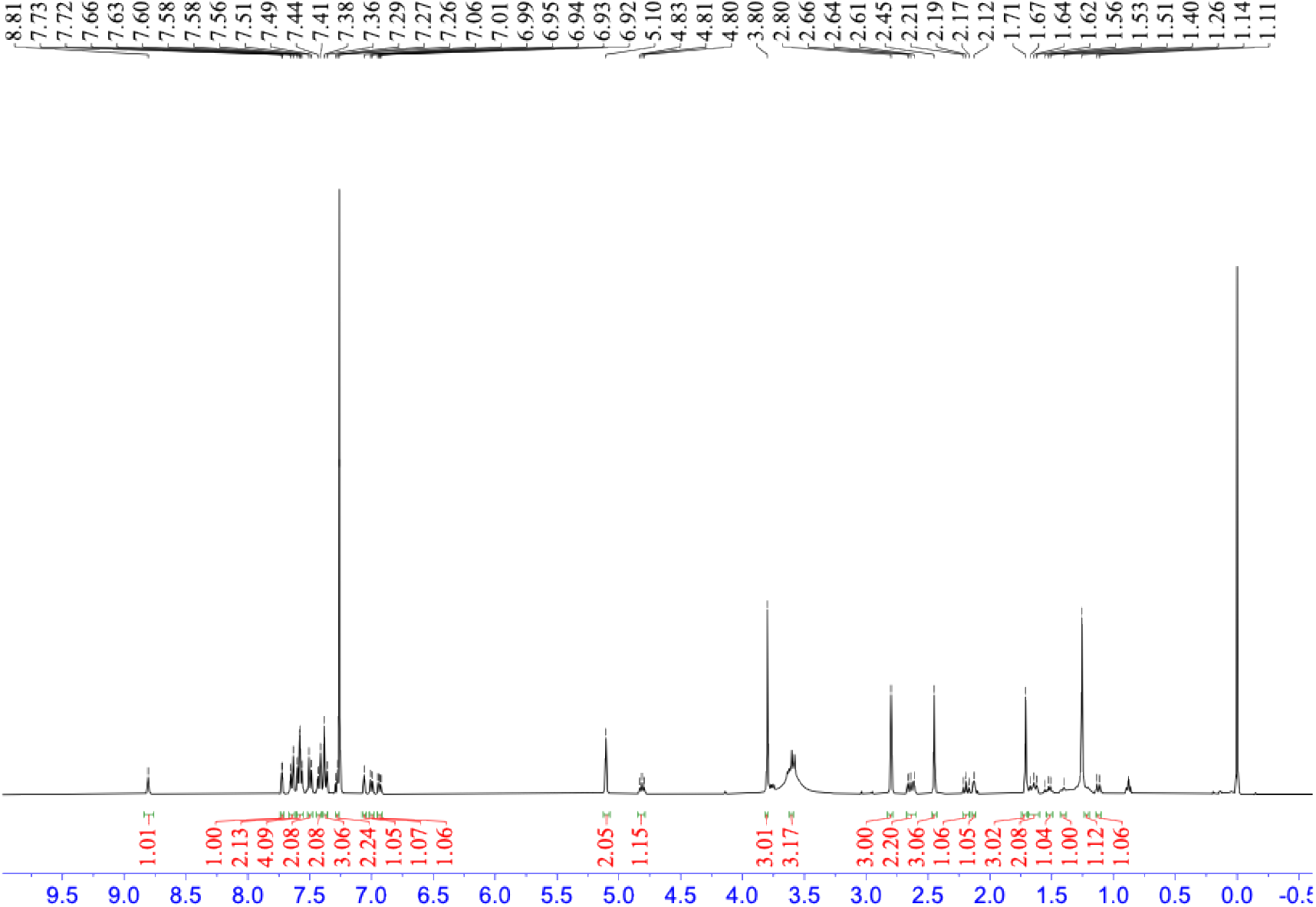

HRMS spectra of **9**

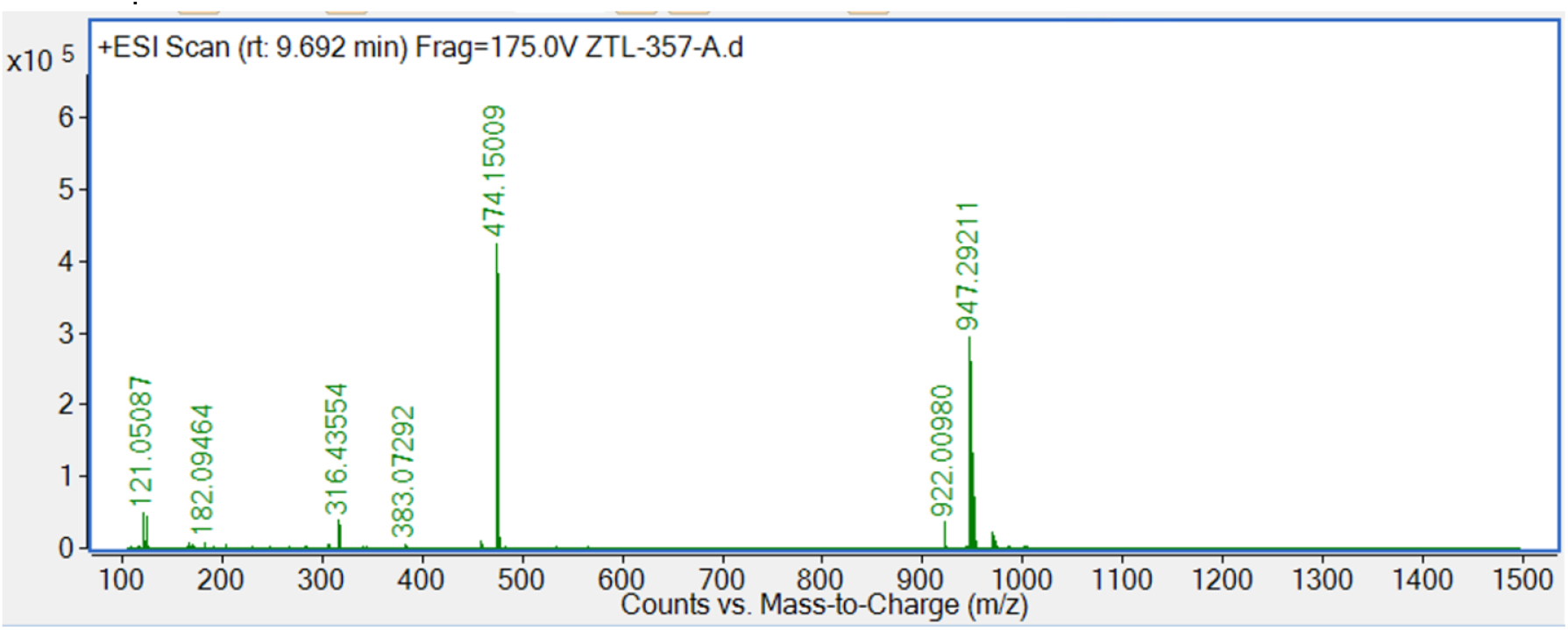

